# Long-range somatic structural variation calling from matched tumor-normal co-assembly graphs

**DOI:** 10.1101/2024.07.29.605160

**Authors:** Megan K. Le, Qian Qin, Heng Li

## Abstract

The accurate identification of somatic structural variants (SVs) is a problem with significant applications to clinical cancer research. Though technologies such as long-read sequencing have facilitated the development of more accurate SV calling methods, existing somatic SV callers still struggle with achieving high precision. In this work, we present colorSV, a long-read-based method for calling long-range SVs by examining the local topology of joint assembly graphs from matched tumor-normal samples. colorSV is the first somatic SV calling method that uses a co-assembly approach, as well as the first SV caller that identifies variants by examining characteristics of the assembly graph itself. We demonstrate near-perfect precision and sensitivity for calling translocations on the COLO829 cell line, outperforming four existing somatic SV callers (Severus, Sniffles2, nanomonsv, and SAVANA) in both metrics. We also evaluated colorSV for calling translocations on the HCC1395 cell line, finding that our method achieved a good balance between sensitivity and precision (where the sensitivity was only outperformed by Severus, and the precision was only outperformed by nanomonsv). Our work establishes a novel joint assembly-based strategy for characterizing long-range somatic variation, which could be further expanded or modified for the identification of SVs of different types and sizes.

## Introduction

Structural variations (SVs), which are generally defined as the rearrangement of segments of the genome at least 50 bp long, have been shown to play an important role in cancer development^1–3^. These include large-scale genomic events that exchange regions between separate chromosomes and may induce the emergence of cancerous lesions^4^. Such events may arise either within the germline (as inherited mutations), or as somatic mutations (non-hereditary mutations), requiring the development of specialized methods to detect variants of each type. However, many previous studies have focused on the detection of germline SVs and struggle with the identification of low-frequency somatic variants^5^. Thus, further work toward developing specialized somatic SV calling methods is still needed.

One technological advancement that has facilitated improvements in the accuracy of identification methods is long-read sequencing, which improves the ability to resolve complex regions and rearrangements of the genome. Previous studies have demonstrated improved SV calling performance from long reads compared to short reads^6,7^. However, existing somatic SV calling methods still struggle with achieving high accuracy, particularly when evaluated on precision, with some studies finding that the majority of SVs in merged call sets are caller-specific^8^.

Currently, these existing long-read somatic SV callers identify variants by analyzing reads from both tumor samples and matched normal samples. Most methods are alignment-based, in which raw sequencing reads are aligned to a reference and then compared between the tumor and normal samples. Examples of these methods include Sniffles2^9^, nanomonsv^10^, Severus^11^, DeBreak^12^, pbsv^13^, and cuteSV^14^.

In contrast, some germline SV callers employ an assembly-based strategy, where reads are first assembled into contigs before being aligned to a reference. One study found that for calling germline SVs, long-read assembly-based methods tend to reach higher sensitivity and precision compared to alignment-based methods^15^, since assembled contigs may provide better resolution for large repetitive loci and higher base-level accuracy. One possible avenue for improving somatic SV calling methods therefore lies within the exploration of novel assembly-based approaches.

In particular, co-assembly-based approaches—in which reads from multiple samples are combined to create a single joint assembly—have been previously employed for *de novo* copy number variation detection between two genomes^16^, as well as for calling SVs within microbiomes^17^. Another study demonstrated improved performance for both read alignment and SV calling by using a personalized reference genome assembled from tumor and matched normal samples^18^. Furthermore, a matched tumor-normal comparative assembly approach has been previously employed for single nucleotide variant calling^19^. However, a co-assembly-based approach has not yet been used during the process of somatic SV calling, nor has any method performed SV calling by examining the structure of the assembly graph itself.

In this work, we demonstrate the utility of matched tumor-normal co-assembly graphs for long-range somatic SV detection by characterizing differences in graph topology between true somatic breakpoints and false positives. We introduce colorSV, a long-range SV caller that achieves near-perfect sensitivity and precision for calling translocations on the COLO829 cell line by examining co-assembly graph topology. We additionally demonstrate high precision and improved sensitivity on the HCC1395 cell line compared to published long-read SV callers (Sniffles2 and nanomonsv), as well as comparable performance to unpublished callers (Severus and SAVANA). Our results provide a novel joint assembly approach toward somatic SV calling that can be further expanded for the identification of additional types of variation.

## Results

### Overview of colorSV method

For each dataset, we first matched the cancer cell line to a lymphoblastoid cell line from the same patient, then jointly assembled the reads from samples of both cell lines to obtain a single co-assembly graph (**Figure 1**). Next, we identified unitigs in the co-assembly graph that were only supported by reads from the tumor samples. We further filtered this set of tumor-only unitigs to exclude those with only 1 supporting read and those that contained telomere sequences. We then mapped the final set of tumor-only unitigs to the T2T-CHM13 reference genome^20^. To generate a set of potential somatic breakpoints, we identified the unitigs that mapped as split alignments and marked these as our initial candidates.

**Figure 1.**
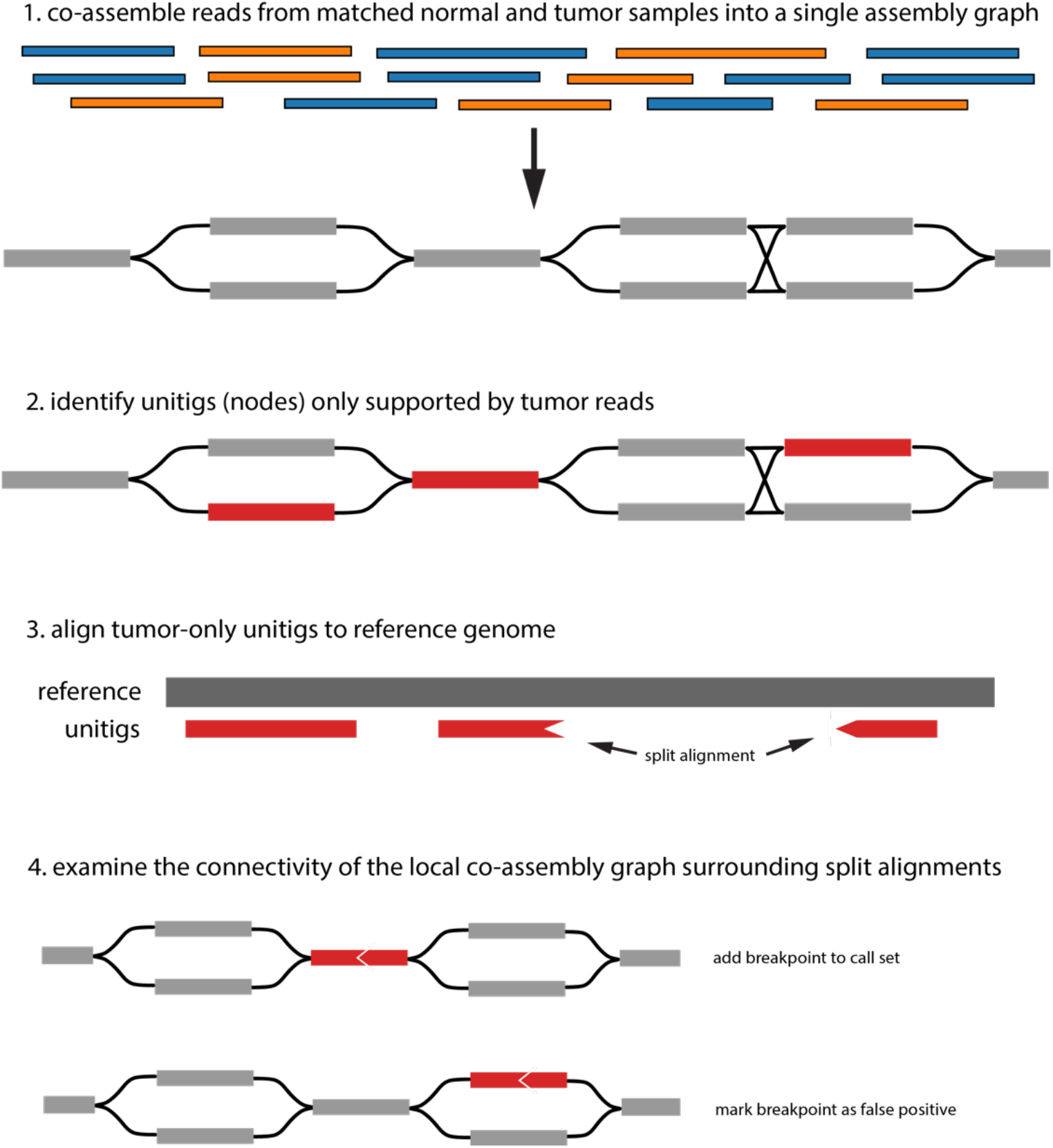
Overview of the colorSV SV calling process. First, reads from matched normal and tumor samples are co-assembled. Next, tumor-only unitigs from the co-assembly graph are aligned to a reference genome, and split alignments are marked as candidate breakpoints. Finally, colorSV only reports breakpoints for which the co-assembly graph is locally disconnected after removing tumor-only unitigs.

To distinguish true somatic breakpoints from germline variation and other false positives, we examined the local topology of the co-assembly graph surrounding the candidate unitigs. Notably, the false positive candidate unitigs tended to be characterized by a bubble-like graph topology, where a parallel path of non-tumor-only unitigs would also connect the candidate’s neighbors (**Figure 2**). On the other hand, true somatic breakpoints tended to connect two subgraphs that were otherwise locally disconnected (**Figure 2**), where the subgraphs contained unitigs corresponding to distant regions of the genome (e.g., separate chromosomes).

**Figure 2.**
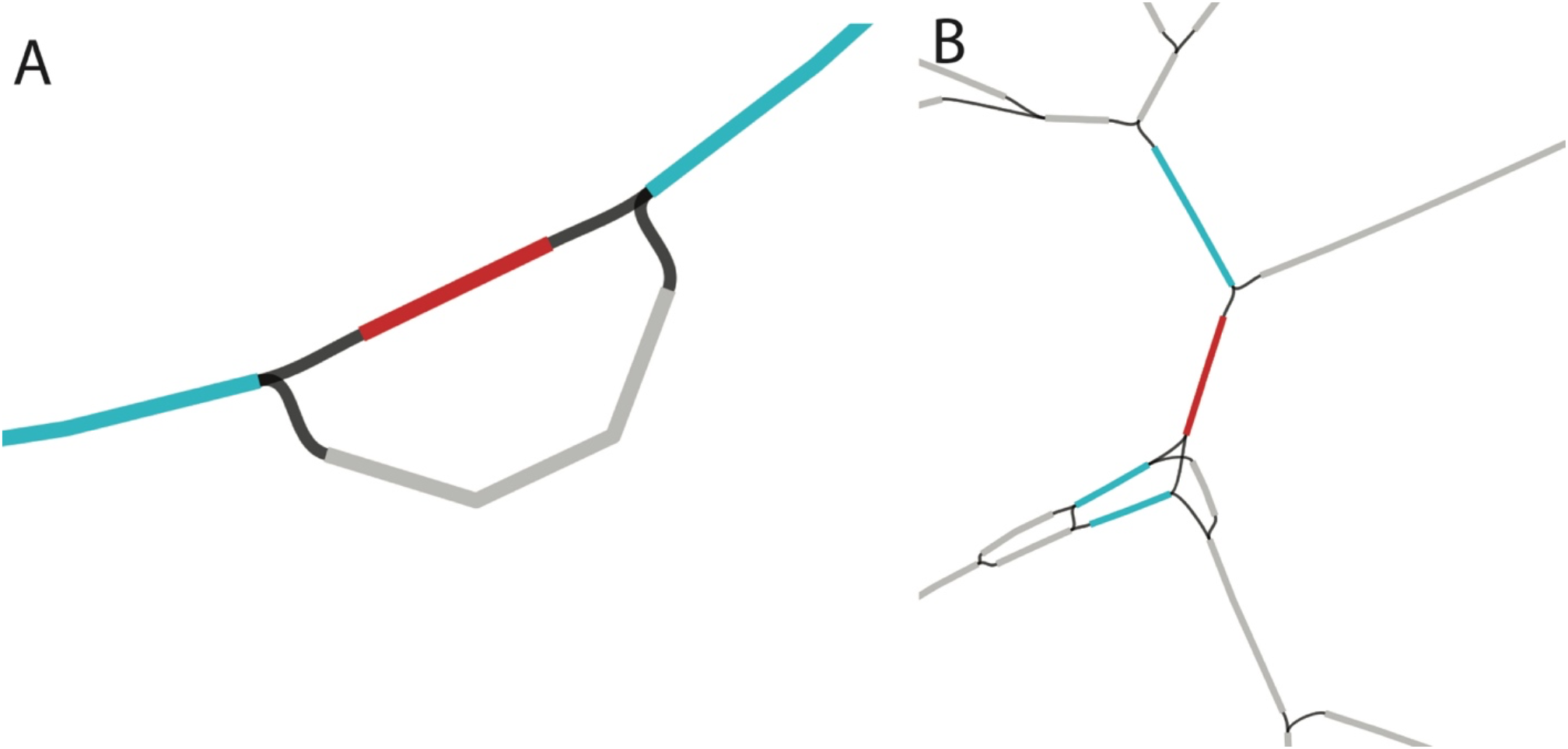
Local co-assembly graph topologies of (**A**) false positive and (**B**) true positive candidate unitigs viewed in Bandage^21^. Candidate unitigs are highlighted in red, while the candidate unitigs’ neighbors are highlighted in blue.

In order to reduce our initial set of candidates to our final call set, we therefore performed checks for the local connectivity of the co-assembly graph surrounding each candidate unitig. For each candidate unitig, we selected a random neighbor and checked whether the other neighbors were reachable within 10 layers of a breadth-first search after removing tumor-only unitigs from the graph. If the candidate’s neighbors were still connected to each other, we marked the candidate as a false positive and removed it from the final call set. Candidates with less than 2 neighbors were also marked as false positives.

Our call set comprised the breakpoints found in the remaining candidate unitigs, which we further filtered by removing breakpoints that occurred within 1Mb of a centromere, as well as unitigs that had an alignment with a mapping quality less than 15.

### Evaluation on the COLO829 cell line

We first tested our method on the COLO829 cell line, a fibroblast cell line derived from a metastasis of a cutaneous melanoma patient. We matched this cell line to the COLO829BL cell line, a B lymphoblast cell line isolated from the peripheral blood of the same subject.

colorSV identified 11 somatic interchromosomal events, all of which appeared to be real based on manual inspection of the reads using the Integrative Genomics Viewer (IGV)^22^. We also compared our results to the 13 translocations found in the reference call set generated by Valle-Inclan et al.^23^, which was curated and experimentally validated using multiple different sequencing technologies. From the reference set, two of the translocations (between chromosomes 6 and 15) were in close proximity to each other and not present in the reads, suggesting that these events are missing due to differences in sample lineage between our data and the reference set samples (**Supplementary Figure 1**).

colorSV identified 10 out of the remaining 11 reference set translocations. This included a translocation between chromosomes 1 and 10 that is 67 bp away from a 2 Mb deletion on chromosome 10 (**Supplementary Figure 2**). However, colorSV called the deletion breakpoint as one of the translocation breakpoints, since the portion of the tumor-only unitig that aligned to the 67 bp region had a MAPQ score that fell below the threshold that was used when extracting SV breakpoints from aligned unitigs.

We did not identify one reference set translocation (between chromosomes 3 and 6) due to an error in the co-assembly process, where the assembler discarded the reads containing the breakpoint, resulting in the breakpoint not being reflected in the final co-assembly graph. This removal may be related to another event on chromosome 3 that is about 1 Mb away and includes breakpoints between chromosomes 10 and 12, perhaps leading the assembler to erroneously remove the reads with the chromosome 6 breakpoints during graph simplification.

We compared the results of colorSV to four other SV callers: Sniffles2^9^, nanomonsv^10^, Severus^11^, and SAVANA^24^. When evaluating the performance of other methods, we considered the set of known translocations to be the 11 translocations from the Valle-Inclan et al. reference set, along with the additional translocation between chromosomes 6 and 15 that was identified by colorSV and appears to be real from manual inspection of supporting reads (**Supplementary Figure 3**).

In comparison to our results, the Severus caller also identified 11 translocations, 9 of which were found in the colorSV call set. Of the 2 unique Severus calls, one appeared to be real based on the supporting reads and was also found in the reference set, while the other was believed to be a false positive. The nanomonsv and Sniffles2 callers both successfully identified 3 translocations, ultimately missing 9 known translocations while also reporting the same false positive translocation as Severus. The SAVANA caller correctly identified 6 translocations, missing 6 known translocations while reporting the same false positive translocation as the other callers (**Table 1**).

**Table 1.** Performance of different SV callers with respect to a known set of translocations in the COLO829 cell line (using the T2T-CHM13 reference). Our method achieves perfect precision and an improvement in sensitivity over other methods. Unpublished callers are denoted by an asterisk.

| Method | Sensitivity | Precision |
| --- | --- | --- |
| colorSV | 11 / 12 | 11 / 11 |
| Sniffles2 | 3 / 12 | 3 / 4 |
| nanomonsv | 3 / 12 | 3 / 4 |
| Severus* | 10 / 12 | 10 / 11 |
| SAVANA* | 6 / 12 | 6 / 7 |

All of the other SV callers found a translocation between 6q23.2 and 7q35. The translocation was apparently supported by read alignment against T2T-CHM13 (**Supplementary Figure 4**) but not by alignment against GRCh38. Valle-Inclan et al.^23^, which only looked at alignments to GRCh38, did not report this translocation, either. We note that 7q35 harbors a complex germline segmental duplication around gene ARHGEF35 and OR2A42 [PMID: 25621458], where the COLO829BL haplotypes are distinct from T2T-CHM13 based on the germline-only assembly. When we aligned tumor reads or tumor-only unitigs to the germline-only assembly, we saw evidence of inversions and duplications, suggesting the presence of local somatic SVs, but there was no translocation signal. We speculate that the combination of complex germline SVs and local somatic SVs resulted in incorrect read alignments and subsequently the spurious translocation signal. We did not see the translocation signal in the GRCh38 alignment, most likely because COLO829BL is more similar to GRCh38 than to T2T-CHM13 at this locus. By taking advantage of *de novo* assembly, colorSV is less affected by germline SVs between COLO829 and the reference and does not call this false positive.

colorSV additionally identified a 6q12-15q11.2 translocation that was not found by Valle-Inclan et al.^23^, most likely because chimeric reads at 15q11.2 on T2T-CHM13 were best mapped to KI270727, an unlocalized contig in GRCh38. The other SV callers also missed this SV, possibly due to a long segmental duplication at 15q11.2 that led to lower mapping quality for ∼20kb HiFi reads. colorSV extracted a ∼60kb unitig containing the translocation. The longer flanking sequence lengths helped improve the signal.

We also investigated the performance of our method for other SVs larger than 1 Mb. The Valle-Inclan et al. reference set reported the 2 Mb deletion on chromosome 10 that falls 67 bp away from the translocation to chromosome 1, which were called together by colorSV. Our method also identified a large somatic deletion on chromosome 7 not reported in the reference set, which we believe to be a true event through manual inspection of the aligned supporting reads.

### Evaluation on the HCC1395 cell line

We also tested our method on the HCC1395 and HCC1395BL cell lines, which are matched cancer-lymphoblastoid cell lines from a patient with ductal carcinoma. We identified 101 translocations, of which 89 were also identified by the Severus caller. Of the 12 we identified that were missed by Severus, 9 appeared to be real through manual inspection of the supporting reads with IGV, while 3 were believed to be false positives.

Severus additionally identified 27 events that we did not identify. Of these 27, 5 fell within centromere regions, and 3 others were believed to be false positives (**Supplementary Figure 5**). The other 19 were believed to be true somatic translocations. When investigating why colorSV was unable to call these translocations, we found that colorSV incorrectly classified 7 candidates due to the topology search criteria (including 3 cases where the candidate unitig only had 1 neighboring unitig in the co-assembly graph). The other 12 translocations were unable to be recovered because the assembler either removed the reads containing the breakpoints from the assembly graph, or the reads with the breakpoints were collapsed into unitigs that contained reads from the normal sample and represented sequences that did not reflect the somatic variation.

We also calculated proxies for the false negative and false discovery rates of each of the five SV callers (colorSV, Sniffles2, nanomonsv, Severus, and SAVANA) by generating a reference call set from the results of the other methods^11^. For each SV caller, we evaluated the false discovery rate by using the union of the other SV call sets as a reference set. To evaluate the false negative rate, we generated reference sets that consisted of variants reported by at least two other callers. We found that colorSV had the second best performance for both metrics, with only Severus achieving a lower false negative rate and only nanomonsv achieving a lower false discovery rate (**Figure 3**, **Table 2**). We caveat that since colorSV employs a co-assembly-based approach while the other SV callers use read-based approaches, the errors within the call sets of the other methods are likely to be more similar to each other compared to the errors generated by colorSV. This means that we may expect a relative decrease in performance by colorSV on these metrics.

**Figure 3.**
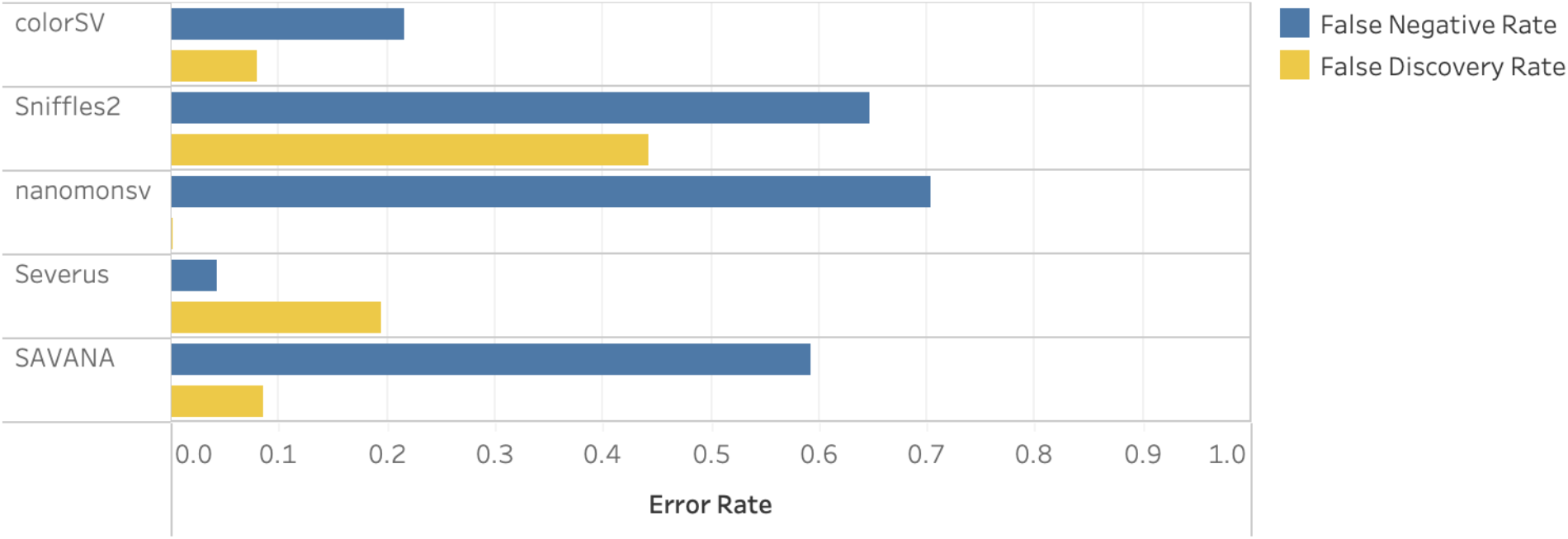
Estimated false negative and false discovery rates for each evaluated SV caller. The reference sets used to calculate the false negative rates consisted of variants reported by at least two other methods. The reference sets used to calculate the false discovery rates were generated by taking the union of the other methods’ call sets.

**Table 2.** Estimated false negative and false discovery rates for each evaluated SV caller. The reference sets used to calculate the false negative rates consisted of variants reported by at least two other methods. The reference sets used to calculate the false discovery rates were generated by taking the union of the other methods’ call sets.

|  | Sensitivity |  |  | Precision |  |  |
| --- | --- | --- | --- | --- | --- | --- |
|  | Total Reference Set Size | Unique Calls in Reference Set | False Negative Rate | Total Call Set Size | Unique Calls in Call Set | False Discovery Rate |
| **colorSV** | 60 | 13 | 0.2167 | 101 | 8 | 0.0792 |
| **Sniffles2** | 85 | 55 | 0.6471 | 79 | 35 | 0.443 |
| **nanomonsv** | 91 | 64 | 0.7033 | 28 | 0 | 0 |
| **Severus** | 47 | 2 | 0.0426 | 123 | 24 | 0.1951 |
| **SAVANA** | 81 | 48 | 0.5926 | 47 | 4 | 0.0851 |

We also performed a pairwise comparison of the overlap between the call sets generated by different callers, which can be found in **Supplementary Table 1**.

## Discussion

Our method demonstrates improved precision and sensitivity over state-of-the-art published methods (Sniffles2^9^, nanomonsv^10^) and unpublished methods (Severus^11^, SAVANA^24^) for calling translocations on the COLO829 cell line, attaining perfect precision and a 11/12 sensitivity. On the HCC1395 cell line, colorSV additionally performed better than most other SV callers using proxies for sensitivity and precision, where the sensitivity was only outperformed by Severus, and the precision was only outperformed by nanomonsv.

By using an approach that leverages information from *de novo* co-assembly, colorSV is less susceptible to errors that may arise as a result of germline SVs. Additionally, the use of unitigs rather than individual reads for performing breakpoint identification may facilitate more accurate alignment and SV detection due to the longer sequence lengths, particularly for somatic events contained within complex or repetitive regions.

One limitation of our approach is its reliance on current assembly tools being able to generate accurate co-assembly graphs, meaning that our method is more likely to fail near complex regions of the genome or where there are clusters of multiple genomic events within a small region (such as in the case of the majority of the known translocations that we missed in the COLO829 and HCC1395 cell lines). However, as assembly methods continue to improve and become more robust for resolving complex regions of the genome, the sensitivity of our framework will improve as well.

In this work, we focused on long-range structural variation (events happening at the scale of 1 Mb) due to the distinctive topology of co-assembly graphs around true somatic breakpoints. The same topology search would not work for short-range structural variants, since the local co-assembly graph for most short-range events would remain connected even without tumor-only unitigs due to the close proximity of neighboring unitigs on the genome. However, it is possible that a similar examination of co-assembly graphs using different search parameters or criteria may be able to identify somatic structural variations of different types or scales.

## Materials and methods

### colorSV

The code for our unitig filtering and co-assembly graph topology search is available at https://github.com/mktle/colorSV.

We co-assembled the matched tumor and normal samples with hifiasm v0.19.6^25^, and we performed our analyses on the generated raw unitig graph. We used minimap2 v2.26 with the long-read presets -ax map-hifi and the -s50 flag to map the tumor-only unitigs to the T2T-CHM13 reference genome^20,26^. The evaluation between callsets was performed using minisv^27^.

### Other SV callers

For each SV caller, we used minimap2 v2.28-r1209 with the same alignment parameters as was used in the colorSV pipeline (long-read presets -ax map-hifi and the -s50 flag) and samtools v1.20-3 to prepare the sorted CRAM.

We used Severus (v1.0) in the tumor-normal pair mode to generate the comparison translocation set. First, for the paired normal sample, small germline SNV and INDELs were called from the sorted CRAM file using the long-read WGS variant caller Clair3 with the pre-trained hifi_revio model and haplotype phasing mode option --enable_phasing -- longphase_for_phasing. Second, for both the tumor sample and its paired normal sample, we tagged the haplotype classes and block information for each read using whatshap haplotag, inputting the phased germline VCF and sorted CRAM with the option --ignore-read-groups --tag-supplementary --skip-missing-contigs. Third, we fed the haplotagged tumor and normal reads with the phased germline VCF into Severus to call somatic SVs with the variable number tandem repeat (VNTR) aware mode; default options were used for filtering reads and SVs. Both the Severus source code and VNTRs are available at https://github.com/KolmogorovLab/Severus.

Sniffles2 (v2.2) was not originally designed to identify somatic SVs in a tumor-normal mode. Thus, we adapted its multi-sample workflow for this purpose. First, the sorted CRAM was processed using the Sniffles2 tandem repeat aware mode --tandem-repeats to generate an intermediate snf format output for the tumor and normal samples separately; default options were used for reads filtering. Second, we combined the tumor and normal snf to call the SVs existing in both tumor and normal samples using the default options with --tandem-repeats. Third, we identified somatic SVs using minisv.js snfpair -t 1 -n 2. The Sniffles software and tandem repeat BED files were obtained from https://github.com/fritzsedlazeck/Sniffles.

We ran SAVANA (v1.0.5) with tumor and normal sorted CRAM files using the default options and restricted the SV calls to autosome and gender contigs. Nanomonsv (v0.7.0) was used to benchmark the somatic SVs calls with the tumor and normal sorted CRAM files as processed above.

All of the SV callers were installed using the miniconda virtual environments and developed as an end-to-end computational pipeline based on a Directed Acyclic Graph-based workflow tool Snakemake^28^.

### COLO829 and HCC1395 Datasets

We performed our analyses on PacBio Revio platform datasets. The HiFi reads stored in unaligned uBAM format for HCC1395 and HCC1395BL were downloaded from the PacBio website at https://downloads.pacbcloud.com/public/revio/2023Q2/HCC1395/. The HiFi reads stored in unaligned uBAM format for COLO829 and COLO829BL were downloaded from the PacBio website at https://downloads.pacbcloud.com/public/revio/2023Q2/COLO829/COLO829/.

## Authors’ contributions

H.L. conceived and supervised the study. M.K.L. developed the method and drafted the manuscript. Q.Q. generated comparison call sets from other variant callers.

## Acknowledgments

We thank Yujie Guo and Haoyu Cheng for helpful discussions. Megan Le was supported by an MIT Electrical Engineering and Computer Science Great Educators Fellowship.

## Supplementary materials

**Supplementary Figure 1.**
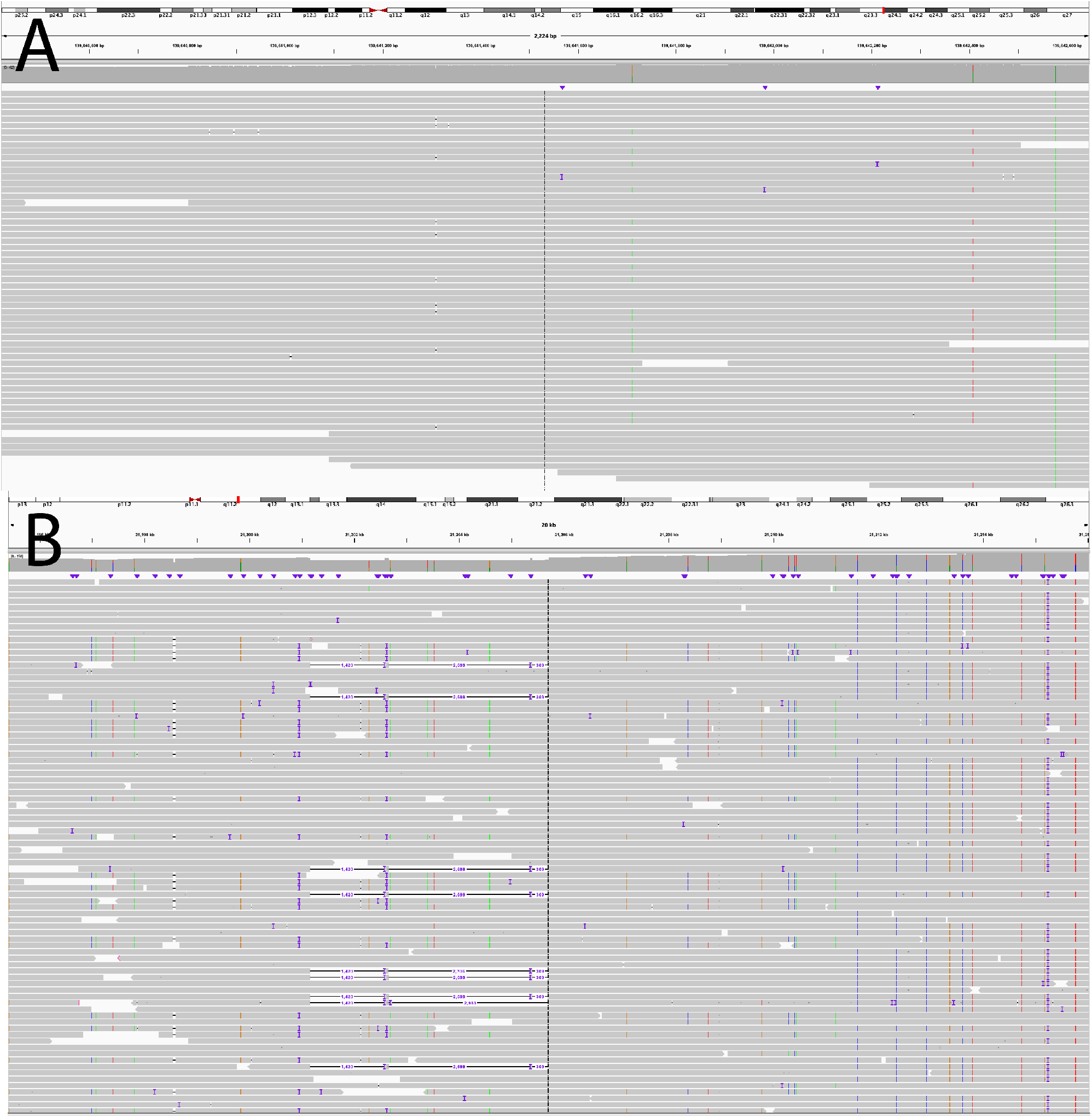
IGV screenshots of the regions on (**A**) chromosome 6 and (**B**) chromosome 7 surrounding the locations of the Valle-Inclan et al.^23^ COLO829 reference set translocations that were believed to not be present in our tumor samples.

**Supplementary Figure 2.**
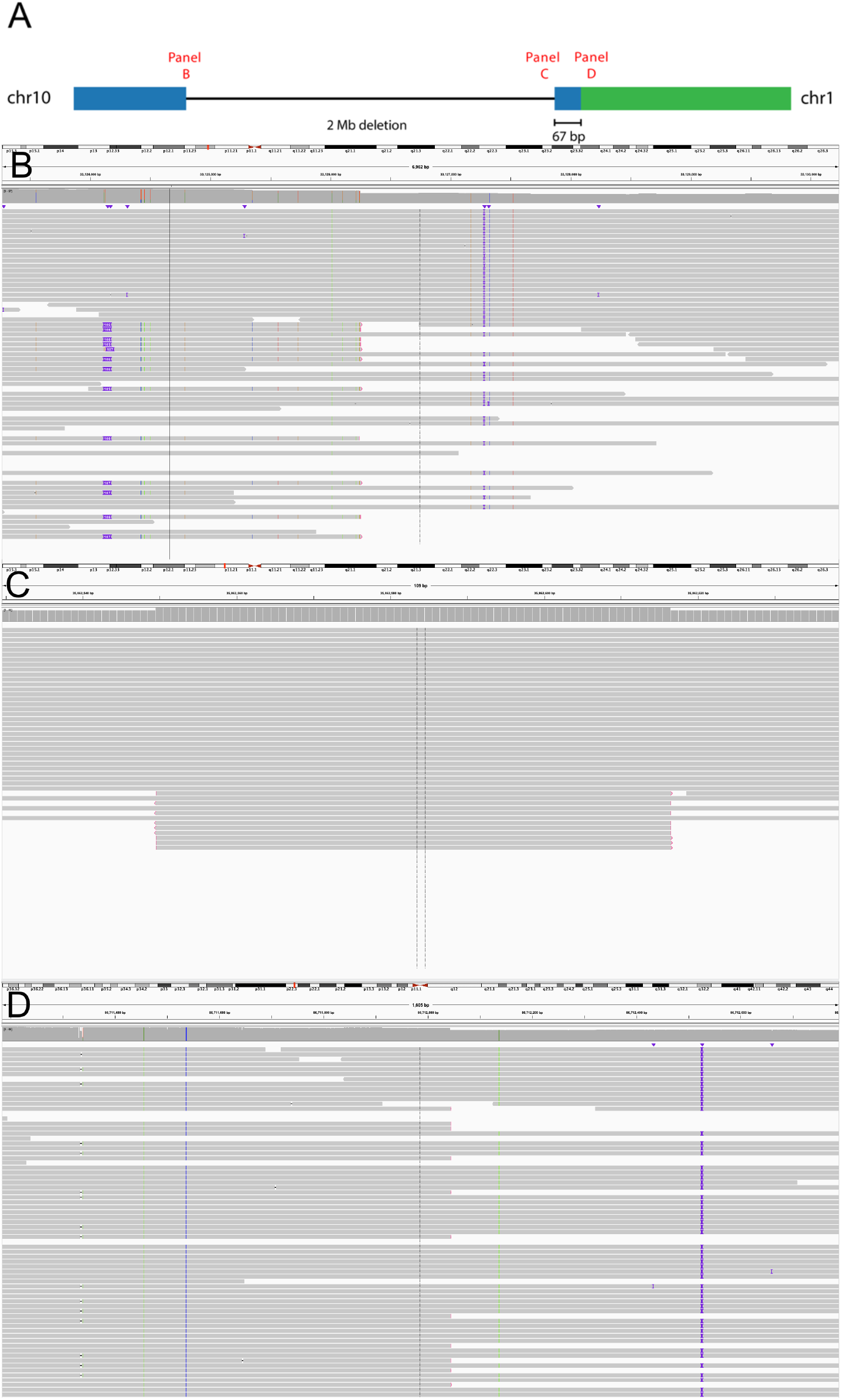
(**A**) Illustration of the somatic deletion and nearby somatic translocation on chromosome 10 in the COLO829 cell line. IGV screenshots of the tumor sample alignments at (**B**) the beginning of the deletion, (**C**) at the 67 bp segment that separates the deletion and translocation, and (**D**) at the breakpoint location on chromosome 1.

**Supplementary Figure 3.**
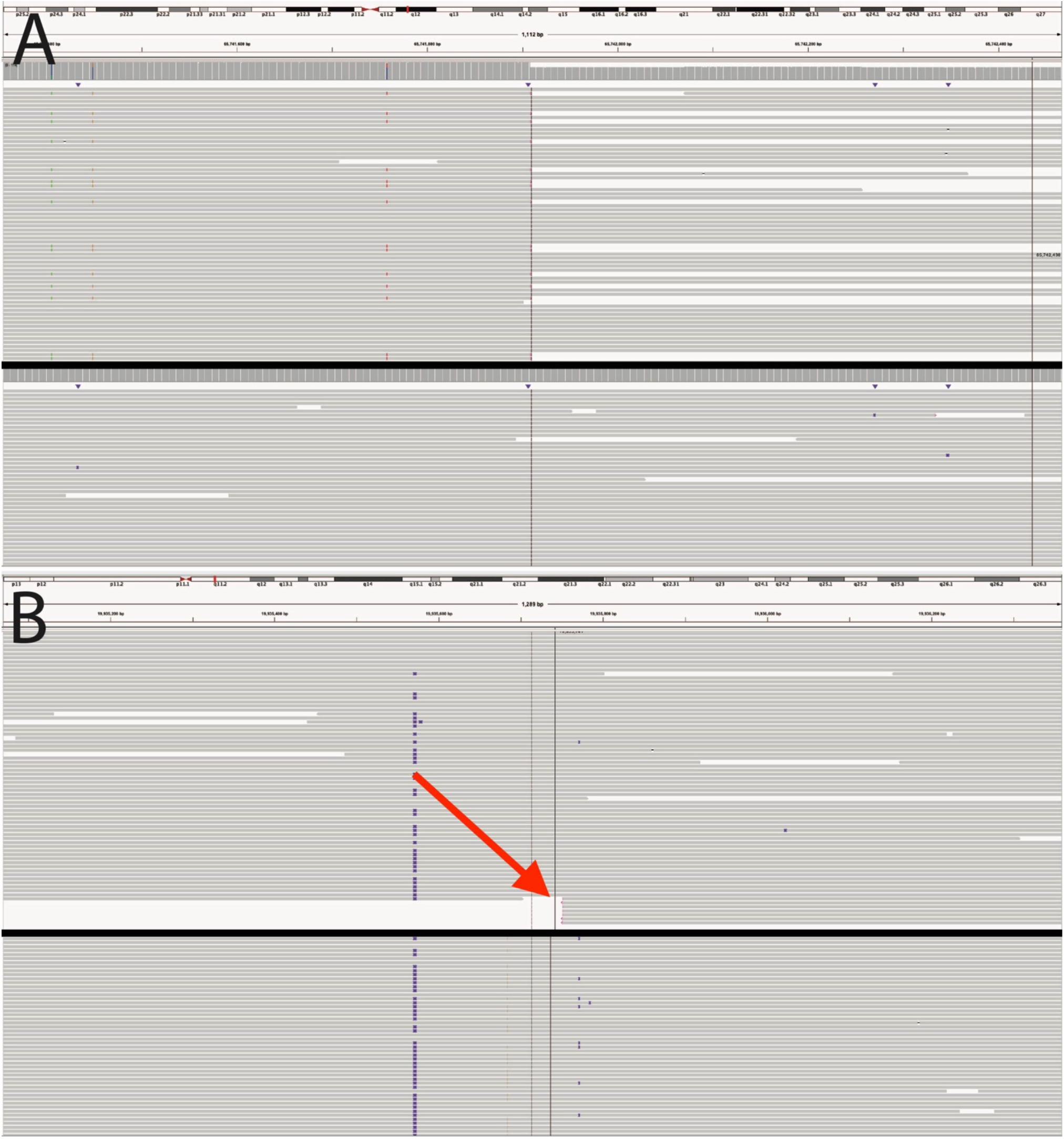
IGV screenshots of the translocation between (**A**) chromosome 6 and (**B**) chromosome 15 that was reported by colorSV and not by other SV callers on the COLO829 cell line. The top track in each panel displays the tumor sample reads, while the bottom track displays the matched normal sample reads. The red arrow indicates the breakpoint on chromosome 15 that is adjacent to the breakpoint on chromosome 6.

**Supplementary Figure 4.**
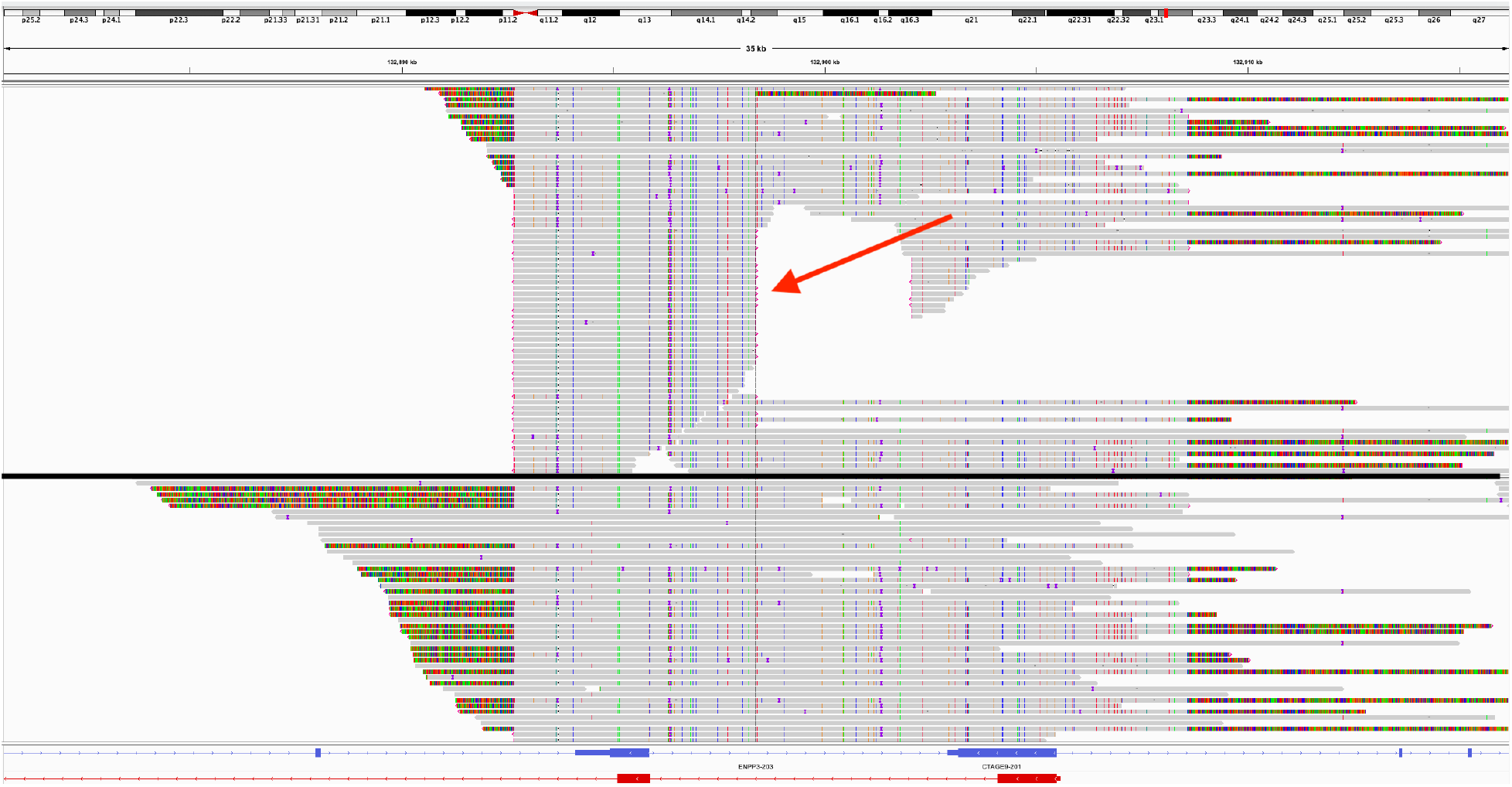
IGV screenshot of the false positive translocation (between chromosomes 6 and 7) called by all other SV callers on the COLO829 cell line, where the top track displays the tumor sample read alignments and the bottom track displays the matched normal sample read alignments on chromosome 6. The red arrow indicates the breakpoint only found in the tumor sample, which results in a false signal for SV callers despite being believed to be an alignment artifact. The clipped reads at the other two sites present in both the tumor and the normal samples also aligned to chromosome 7. When we aligned the reads or tumor-only unitig back to an assembly of the normal sample reads or to the GRCh38 reference, the translocation signal disappeared.

**Supplementary Figure 5.**
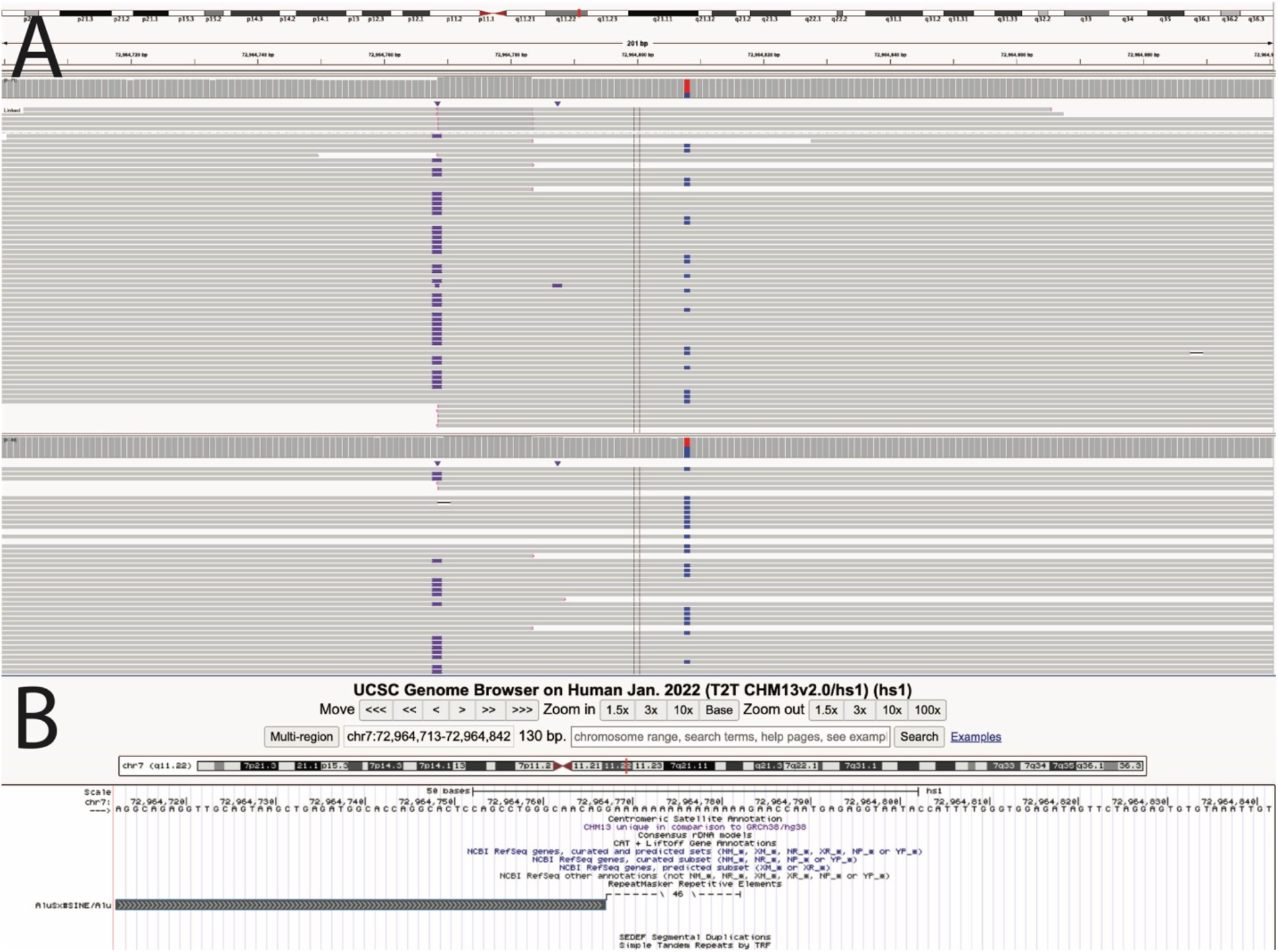
(**A**) IGV screenshots of one of the translocations (between chromosomes 7 and 20) reported by Severus on the HCC1395 cell line, which was believed to be a false positive. The top track displays alignments on chromosome 7 from the tumor sample, while the bottom track displays alignments on chromosome 7 from the matched normal sample. Though some reads have 500 bp supplementary alignments to chromosome 20, others have a 1 kb insertion at the same site (represented by the purple bars). This pattern is present in both the tumor and the normal reads. (**B**) The UCSC Genome Browser indicates the presence of an AluSx repeat at this location, which could be causing a spurious signal due to difficulty performing alignment.

**Supplementary Table 1.** Overlap in translocation call sets on the HCC1395 cell line between our method and other SV callers. The diagonal entries indicate the total number of translocations reported by each caller. Each off-diagonal entry reports the fraction of the row call set that was also found in the column call set. Higher row values are correlated with overall higher sensitivity, while higher column values are correlated with overall higher precision. Unpublished methods are denoted by an asterisk.

|  | colorSV | Sniffles2 | nanomonsv | Severus* | SAVANA* |
| --- | --- | --- | --- | --- | --- |
| colorSV | 101 | 0.4810 | 0.8571 | 0.6992 | 0.7234 |
| Sniffles2 | 0.3267 | 79 | 0.4286 | 0.2846 | 0.4043 |
| nanomonsv | 0.2772 | 0.1772 | 28 | 0.2195 | 0.4043 |
| Severus* | 0.8812 | 0.5316 | 0.9643 | 123 | 0.8511 |
| SAVANA* | 0.3762 | 0.2911 | 0.6786 | 0.3252 | 47 |

